# Behavioral variation correlates with differences in single neuron serotonin receptor subtype expression within and across species

**DOI:** 10.1101/231829

**Authors:** A.N. Tamvacakis, A. Senatore, P.S. Katz

## Abstract

The marine mollusc, *Pleurobranchaea californica* varies daily in whether it swims and this correlates with whether serotonin (5-HT) enhances the strength of synapses made by the swim central pattern generator neuron, C2. Another species, *Tritonia diomedea*, reliably swims and does not vary in serotonergic neuromodulation. A third species, *Hermissenda crassicornis*, never produces this behavior and lacks the neuromodulation. We found that expression of particular 5-HT receptor genes in C2 correlates with swimming. Seven 5-HT receptor subtype genes were identified from whole-brain transcriptomes. We isolated individual C2 neurons and sequenced their RNA or measured 5-HT receptor gene expression using quantitative PCR. C2 neurons isolated from *Pleurobranchaea* individuals that produced a swim motor pattern just prior to isolation expressed the 5-HT2a and 5-HT7 receptor genes, as did the *Tritonia* samples. These subtypes were absent from C2 neurons isolated from *Pleurobranchaea* individuals that did not *s*wim that day and from *Hermissenda* C2 neurons. Expression of other receptors did not correlate with swimming. This suggests that 5-HT2a and 5-HT7 receptors mediate the modulation of C2 synaptic strength and play an important role in swimming. Furthermore, the results suggest that regulation of receptor expression might underlie daily changes in behavior as well as behavioral evolution.

## Introduction

It has been proposed that differences in behavior could arise through differences in neuromodulatory signaling [1–3]. Serotonin (5-HT) alters behavior by modulating neuronal and synaptic properties [4–6]. Many 5-HT receptor subtypes have been identified [7]. Variation in the expression of particular 5-HT receptor subtypes could alter the effect of 5-HT on a neuron.

We examined the expression of genes for 5-HT receptors in single identified neurons of three marine molluscan species (*Tritonia diomedea*, *Hermissenda crassicornis*, and *Pleurobranchaea californica*). These sea slugs have large neurons that can be individually identified from animal to animal within a species, as is common with invertebrates [8, 9]. Furthermore, homologous neurons can be identified across species [10]. Moreover, the roles of these neurons in simple behaviors have been determined.

The nudibranch *Tritonia diomedea* (synonymous with *Tritonia tetraquetra*, Pallas 1888) [11] swims by producing alternating dorsal and ventral (DV) body flexions. The central pattern generator (CPG) underlying the swim motor pattern contains the identified neurons C2 and DSI [12, 13]. *Pleurobranchaea californica* also swims with DV body flexions and has neurons homologous to C2 and DSI in its swim CPG [14–16]. These neurons also can be identified in the nudibranch, *Hermissenda crassicornis*, even though *Hermissenda* lacks this swimming behavior [14, 17].

Serotonergic neuromodulation plays a central role in the production of the swim motor pattern in both *Tritonia* and *Pleurobranchaea*. Application of 5-HT to the isolated *Tritonia* brain is sufficient to evoke a swim motor pattern and injection of the animal with 5-HT evokes a swim [18]. Stimulation of the serotonergic DSI or 5-HT application increases the strength of C2 synapses [19, 20]. In *Tritonia*, this was shown to be caused by neurotransmitter release from C2 through an increase in spike-evoked Ca^2+^ levels in C2 terminals [21]. Furthermore, DSI stimulation affects C2 after-hyperpolarization and elicits both fast and slow excitatory post-synaptic potentials [19, 22, 23]. The 5-HT receptor antagonist, methysergide, blocks the neuromodulatory actions of DSI on C2 synapses [20, 22]. Methysergide also blocks production of the swim motor pattern in the isolated brain and prevents swimming when injected into the intact animal [18].

Unlike in *Tritonia* and *Pleurobranchaea*, neither DSI nor 5-HT modulates C2 synaptic strength in *Hermissenda*. Furthermore, application of 5-HT to the isolated *Hermissenda* brain fails to evoke a swim motor pattern, and injection of *Hermissenda* with 5-HT does not cause the animal to swim [20]. Here, we investigated whether the correlation between swimming behavior and serotonergic modulation of C2 neurons could be caused by species differences in expression of particular 5-HT receptors in C2.

Unlike in *Tritonia*, swimming in *Pleurobranchaea* is not consistent; the same animal varies in its propensity to swim when tested over the course of several days [20]. Similarly, some isolated brain preparations do not exhibit bursting activity typical of a swim motor pattern. It was shown that the extent of serotonergic enhancement of C2 synaptic strength correlated with the number of burst cycles in the motor pattern [20]. Here, we examined expression of 5-HT receptors in *Pleurobranchaea* C2 neurons to determine whether it could underlie the variability in swimming behavior.

## Methods

### Animals

*Tritonia diomedea* specimens were collected by Living Elements LLC (Vancouver, B.C.). *Hermissenda crassicornis* and *Pleurobranchaea californica* specimens were collected by Monterey Abalone Co. (Monterey, CA). All animals were housed at 10°C in recirculating artificial seawater (ASW, Instant Ocean). Individual *Tritonia* were anaesthetized before dissection using cold temperature and the other species were anaesthetized by injecting 0.33 M magnesium chloride into the body cavity.

### Whole-brain RNA extraction, cDNA production, and transcriptome sequencing

For whole-brain RNA extraction, brains were dissected from the animal, cleaned of connective tissue, flash-frozen using liquid nitrogen, and then stored at -80°C. RNA extraction was performed using the RNeasy Plus Universal Mini Kit (Qiagen). RNA extracts were quantified using Nanodrop (Thermo Fisher). RNA was reverse transcribed to cDNA using Superscript IV (Thermo Fisher) following manufacturer’s instructions.

For the *Pleurobranchaea* whole brain transcriptome, additional RNA was extracted as described above. RNA was reverse transcribed, and sequenced on an Illumina Hi-Seq2500, and assembled as described previously for *Tritonia* and *Hermissenda* [24–26]. Metrics for the *Pleurobranchaea* CNS shotgun transcriptome assembly are provided in Supplemental Table 1.

### 5-HT receptor plasmid cloning

Species-specific primers were designed using transcriptome-derived 5-HT receptor sequences from *Tritonia* [24], *Hermissenda* [25], and *Pleurobranchaea* whole brain transcriptomes TBA). The primers (Eurofins) are listed in Supplemental Table 2. PCR was performed to amplify putative 5-HT receptor genes from whole-brain cDNA from each species using *Taq* polymerase (Thermo Fisher). PCR products were gel purified using the Qiaquick Gel Extraction Kit (Qiagen). Resulting DNA was ligated using T4 DNA ligase and inserted into JM109 competent cells (Promega). Cloned plasmids were extracted using the GenElute Plasmid Mini-prep Kit (Sigma Aldrich). Plasmids were sequenced on a 3730 DNA Analyzer (Thermo Fisher), and sequences were aligned against transcriptome sequences using MUSCLE [27] to verify gene identity.

### Synthetic RNA production for quantitative PCR standards

Plasmid DNA sequences for each species-specific 5-HT receptor subtype were linearized with T7- or Sp6-oriented enzyme digests (New England Biosciences) and gel-verified. Digested plasmids were purified using phenol-chloroform extraction and alcohol precipitation, and quantified using a Nanodrop 2000C (Thermo Fisher). Synthetic RNA was produced using either T7 or Sp6 MegaScript Kit (Thermo Fisher) and purified. RNA was quantified using Nanodrop. The copy number for each 5-HT receptor synthetic RNA sample was calculated using the gene-specific molecular weight and RNA concentration [28]. RNA standards were then serially diluted and individually reverse transcribed to cDNA using SuperScript IV (Thermo Fisher).

### Single-neuron isolation

Following initial dissection, the brain, which consists of the fused cerebral, pleural, and pedal ganglia, was placed in either artificial sea water (Instant Ocean) or normal saline (NS) (420 mM NaCl, 10 mM KCl, 10 mM CaCl2, 50 mM MgCl2, 11 mM d-glucose, and 10 mM HEPES, pH 7.5). The connective sheath was removed using fine scissors and forceps.

Intracellular recordings were made using 10-30 MΩ glass microelectrodes connected to an IX2-700 Intracellular Amplifier (Dagan). Extracellular recordings and stimulation of body wall nerves were made by placing them in polyethylene tubing connected to a Model 1700 differential amplifier (A-M Systems). Signals were digitized using the Micro1401 and Spike2 software (Cambridge Electronic Design Limited).

C2 is identified in each species by its white soma in the cerebral ganglion, which is immunoreactive to the neuropeptides FMRFamide and Small Cardioactive Peptide (SCP), and its contralaterally-projecting axon [14, 29, 30]. To electrophysiologically identify C2, neurons were recorded intracellularly and a swim motor pattern was evoked by stimulating a body wall nerve with 20–35 V at 5 Hz for 3 s. In *Pleurobranchaea*, C2 neurons were isolated from bursting and non-bursting preparations; individual preparations were defined as “swimming” if two or more bursts could be evoked, and “non-swimming” if one or no bursts were elicited. Electrical coupling to the contralateral white cell was tested in some *Hermissenda* and non-swimming *Pleurobranchaea* preparations when participation in the swim motor pattern could not be used to help identify C2.

After identification of C2, the brain was bathed in 0.2% Protease IX for five minutes and washed with ASW or NS. For neurons that were to be sequenced, two additional washes of filtered ASW were performed. Individual C2 neurons were removed with forceps and a suction pipette.

After removal from the brain, single C2 neurons were placed in individual 1.5 mL tubes containing distilled water and RNaseOut (Thermo Fisher), and visually inspected under a light microscope to verify that cells were present, intact, and were the only cells in the tube. C2 neurons were discarded if they appeared damaged at any point during isolation, or did not meet species-specific physiological hallmarks of C2 neurons. Neurons were not isolated if the central ganglia appeared damaged. C2 neurons that appeared to have other neurons attached to them following initial isolation were also discarded.

### Single-cell transcriptome assembly and gene identification

For *Tritonia* and *Hermissenda*, five individual C2 cell samples were prepared. One swimming and one non-swimming *Pleurobranchaea* C2 cell sample was prepared. Single cell cDNA was synthesized using the Clontech SmartSeq v4 Ultra-Low Input RNA Kit (Takara). Libraries were prepared and indexed using the Nextera XT DNA Library Preparation Kit and 96-Sample Index Kit (Illumina). Qubit (Thermo Fisher) and Bioanalyzer (Agilent) analyses were used to affirm quantity and quality of cDNA during both protocols. cDNA samples were sequenced on a Hi-Seq2500 (Illumina) with a read depth of 10 million reads per sample. Sequencing reads as fastq files were analyzed using FastQC (Andrews 2010) via BaseSpace analysis (Illumina). For *Tritonia* and *Hermissenda*, the single sample raw reads were concatenated before assembly. All files were then assembled with Trinity [26]. BLAST+ (version 2.4.0) [31] was used to identify genes with queries from whole-brain transcriptomes of the same species. NCBI tBLASTx and MUSCLE alignment [32] were used to confirm phylogenetic identity of select identified genes. The resulting transcriptomes are published as NCBI SRA TBA.

### Single-Neuron cDNA synthesis for quantitative PCR

Neurons used for qPCR were isolated as described above and were transcribed to cDNA using Superscript IV (Thermo Fisher). C2 neuron samples were collected independently for each qPCR trial. Following isolation, the cells were added to a mixture of distilled water and RNaseOut and frozen at -80°C. A master mix was created following the Superscript IV manufacturer’s protocol, and aliquots were added directly to the RNA. The RNA was annealed and cooled on ice. The annealed RNA was then divided by volume equally into two tubes. One tube received reverse transcriptase enzyme plus additional Superscript IV reagents (+RT). The other tube lacked reverse transcriptase (-RT) and received a volume of distilled water in place of the enzyme volume. The RNA samples were then reverse-transcribed following manufacturer’s instructions.

### mRNA quantification using single-neuron and whole brain absolute qPCR

Absolute qPCR was performed for individual genes from each species. qPCR-specific primers were designed against each gene orthologue (Supplemental Table 3). For 5-HT2a in *Tritonia* and *Hermissenda*, two sets of primers were used to measure mRNA expression at multiple locations along the 5-HT2a mRNA. RNA standards were prepared as described above, and were run alongside +RT cell cDNA, -RT samples, and without-template control samples on every qPCR trial. C2 samples were run either as single cells, or pooled cells from multiple animals; raw copy numbers were divided by starting cell amount. Whole brain tissue was run with 150 ng RNA calculated per one triplicate sample. All C2 and whole brain samples, standards, and controls were run in triplicate using Perfecta SYBR Green Supermix with ROX or Low ROX (Quanta Bio) on an Applied Biosystems 7500 or StepOne Plus qPCR machine.

Absolute mRNA copy number values were calculated using the standard curve generated during each trial. If amplification occurred in the –RT samples, it was subtracted from the +RT samples. Trials were omitted if the standard curve efficiency was calculated to be outside the range of 85 to 110%. A melt curve was run on every trial, and samples were omitted if a double peak was detected, or if samples showed multiple bands in an agarose gel (Sigma Aldrich) following the qPCR trial. mRNA copy numbers were calculated per single cell or per whole brain using SigmaPlot (version 10, Systat Software Inc.).

## Results

### Comparison of C2 5-HT receptor expression in Tritonia and Hermissenda

*Tritonia* reliably produces a swim motor pattern in which C2 is rhythmically active (Fig. 1A). In contrast, *Hermissenda* C2 does not exhibit rhythmic bursting in response to body wall nerve stimulation (Fig. 1B).

**Figure 1:**
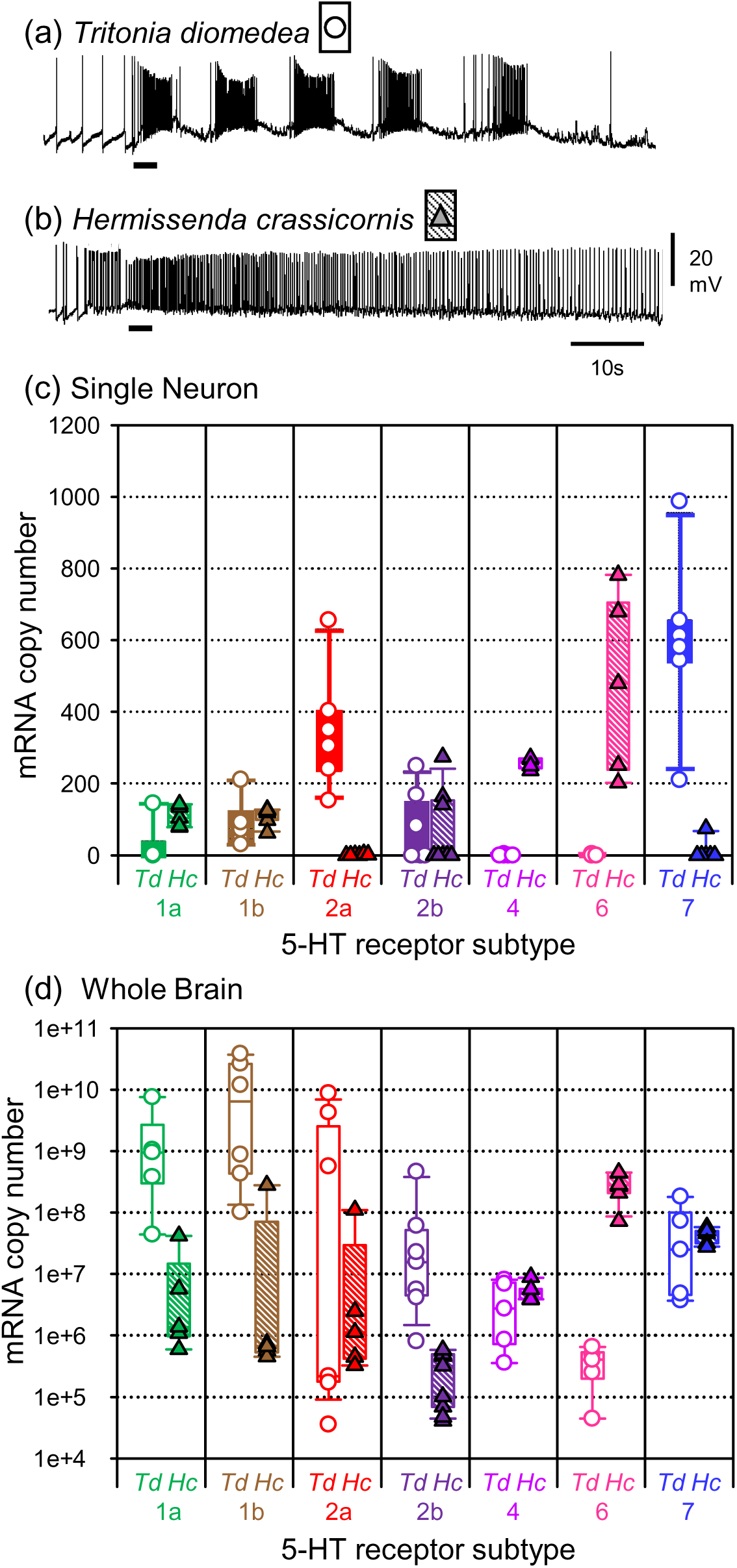
Differences between *Tritonia* and *Hermissenda*. Intracellular recordings of C2 neurons in response to body wall nerve stimulation (black bar) from *Tritonia* (a) and *Hermissenda* (b). Only *Tritonia* exhibits a bursting pattern of action potentials indicative of a swim motor pattern. (c) Single neuron absolute qPCR measurements of mRNA copy number for seven 5-HT receptor genes in *Tritonia* (open circles, white bars) and *Hermissenda* (filled triangles, hatched bars). Symbols represent results from individual C2 neurons. Note that only *Tritonia* expresses 5-HT2a and 5-HT7 receptor genes and that only *Hermissenda* expresses 5-HT4 and 5-HT6 receptor genes. (d) RNA copy number for whole brain tissue. Note that the relative expression of receptors in the whole brain does not correspond to relative expression at the single neuron level with the exception of 5-HT6. For (c) and (d), the box plots represent the range from 25% to 75% of sample expression levels. 95% confidence is represented with error bars.

We examined 5-HT receptor subtype gene expression from single C2 neurons by performing snRNA-seq. Five of the seven receptor subtypes previously identified from whole brain transcriptomes were expressed in the *Tritonia* C2 neuron concatenated transcriptome. These were receptors 5-HT1a, 1b, 2a, 2b, and 7 (Table 1). 5-HT 4 and 5-HT6 were not present in the transcriptome. Thus, only a subset of receptor subtypes were present in *Tritonia* C2 neurons according to snRNA-seq. The C2 transcriptome in *Hermissenda* differed from that of *Tritonia*. It contained 5-HT1a, 1b, 4 and 6 receptors (Table 1).

**Table 1:** Comp IDs of 5-HT receptors in C2 transcriptome assemblies.

| **Receptor** | ***Tritonia*** | ***Hermisenda*** |
| --- | --- | --- |
| 5-HT1a | DN44848_c3_g1_i3 | DN63691_c1_g1_i1 |
| 5-HT1b | DN48323_c0_g1_i1 | DN103990_c0_g1_i1 |
| 5-HT2a | DN61866_c0_g1_i1 | not found |
| 5-HT2b | DN24124_c0_g1_i1 | DN122702_c0_g1_i1 |
| 5-HT4 | not found | DN86843_c0_g1_i1 |
| 5-HT6 | not found | DN57740_c0_g1_i1 |
| 5-HT7 | DN1861_c0_g1_i1 | not found |

The C2 single neuron transcriptome results were independently validated using single neuron qPCR (Fig. 1C). In *Tritonia*, 5-HT2a and 5-HT7 receptor subtypes both had median expression well above 200 copies per cell (Figure 1C). Within these two data sets, variability in copy number across samples was high, with 5-HT2a showing a range between 152 and 655 copies, and 5-HT7 ranging between 209 and 987 copies. 5-HT4 and 5-HT6 were not detected in *Tritonia* C2 neurons, consistent with the RNA-seq results.

Single neuron qPCR of the *Hermissenda* C2 neurons also validated the transcriptome results. With the exception of one *Hermissenda* qPCR sample, which showed low expression of 5-HT7, there was no expression of 5-HT2a or 5-HT7 receptor genes in *Hermissenda* C2 neurons. In contrast, 5-HT4 and 5-HT6, which were not expressed in *Tritonia* C2 neurons, were found to be expressed at greater than 200 copies per cell in *Hermissenda*. Thus, there were differences in the expression of 5-HT receptors in homologous neurons that also differ in their responses to 5-HT and come from species that differ in behavior.

The remaining receptor genes were expressed at low levels in both species (Fig. 1C). 5-HT1a was identified in the transcriptome. However, qPCR was only able to detect this particular transcript in one out of the five *Tritonia* C2 neurons. 5-HT1b was also found in the *Tritonia* transcriptome and consistently expressed at fewer than 200 copies per cell according to qPCR. In *Hermissenda* C2 samples, 5-HT1a and 5-HT1b were expressed below 200 copies per cell.

The 5-HT2b sequence was found in the single-cell transcriptome of both *Tritonia* and *Hermissenda*, but only three out of six *Tritonia* samples and three out eight *Hermissenda* samples expressed 5-HT2b according to qPCR (Fig. 1C). The similarities in the low-level expression of 5-HT1a, 5-HT1b, and 5-HT2b in both *Tritonia* and *Hermissenda* suggests that these receptors are not responsible for the species-differences in the effect of 5-HT on C2.

The differences in 5-HT receptor expression in C2 neurons of *Tritonia* and *Hermissenda* were not reflected in whole brain expression levels (Fig. 1D). All seven of the 5-HT receptor subtypes were measured in whole brain tissue using qPCR. In general the relative mRNA expression of each receptor did not correlate with the single C2 expression. However, 5-HT6 receptor genes were expressed at a lower level in *Tritonia* brains compared to *Hermissenda* brains.

### 5-HT receptor subtypes were identified in Pleurobranchaea whole-brain tissue

Before testing which 5-HT receptor subtypes were expressed in *Pleurobranchaea* C2 neurons, we generated a *Pleurobranchaea* whole brain transcriptome, and mined it for 5HT receptor sequences using BLAST. Orthologues of all seven molluscan 5-HT receptor subtypes [25, 33] were found (Supplemental Table 4). To confirm their sequence identities, we created consensus sequences of each receptor protein coding sequence via cloning and sequencing of complementary DNA generated from whole-brain tissue.

The *Pleurobranchaea* 5-HT receptor subtype consensus amino acid sequences clustered with other subtypes from each 5-HT receptor family (Fig. 2).

**Figure 2:**
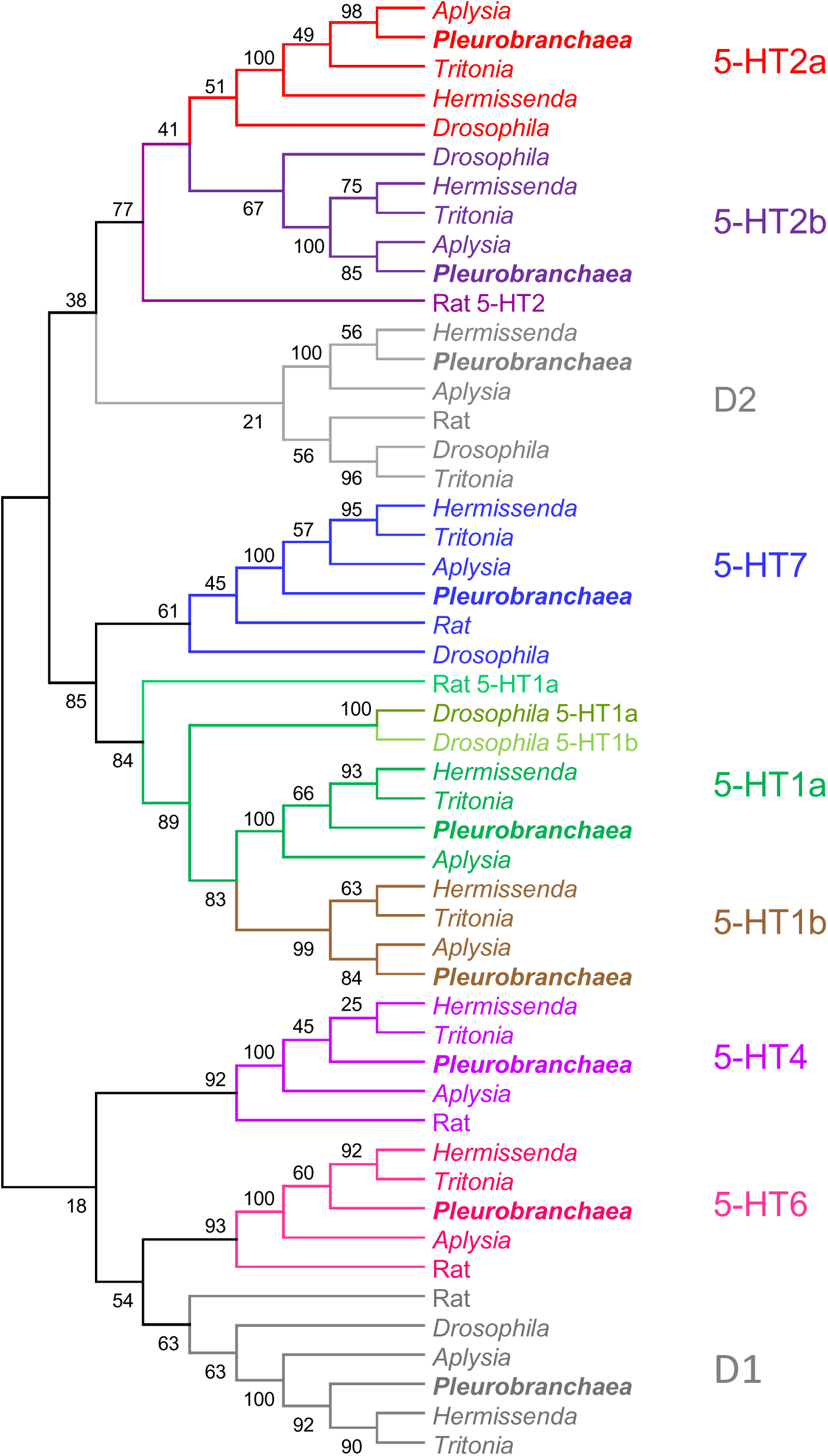
Phylogenetic Tree of 5-HT and Dopamine Receptor Subtypes. A maximum likelihood phylogeny showing seven 5-HT receptor subtypes spanning five families. D1 and D2 dopamine receptors fall within the same ancestral grouping. Previously unpublished *Pleurobranchaea californica* 5-HT receptor sequences were identified from a whole brain transcriptome and sequences were confirmed by plasmid cloning. MUSCLE alignment of predicted amino acids, followed by maximum likelihood tree construction with bootstrapping was done using MEGA6 software [53] . Bootstrap values represent percentage of predicted replicates at each node. Receptor subunits were aligned using conserved transmembrane domain regions.

### Receptor expression differed in swimming and non-swimming Pleurobranchaea

Since *Pleurobranchaea* does not reliably swim, we partitioned samples from this species based on the motor pattern produced at the time that the C2 sample was taken. If the motor pattern consisted of more than a single burst, it was considered as “swimming” (Figure 3a). On the other hand, if stimulation of the body wall nerve produced one or fewer bursts, the individual was categorized as “non-swimming” (Figure 3b).

**Figure 3:**
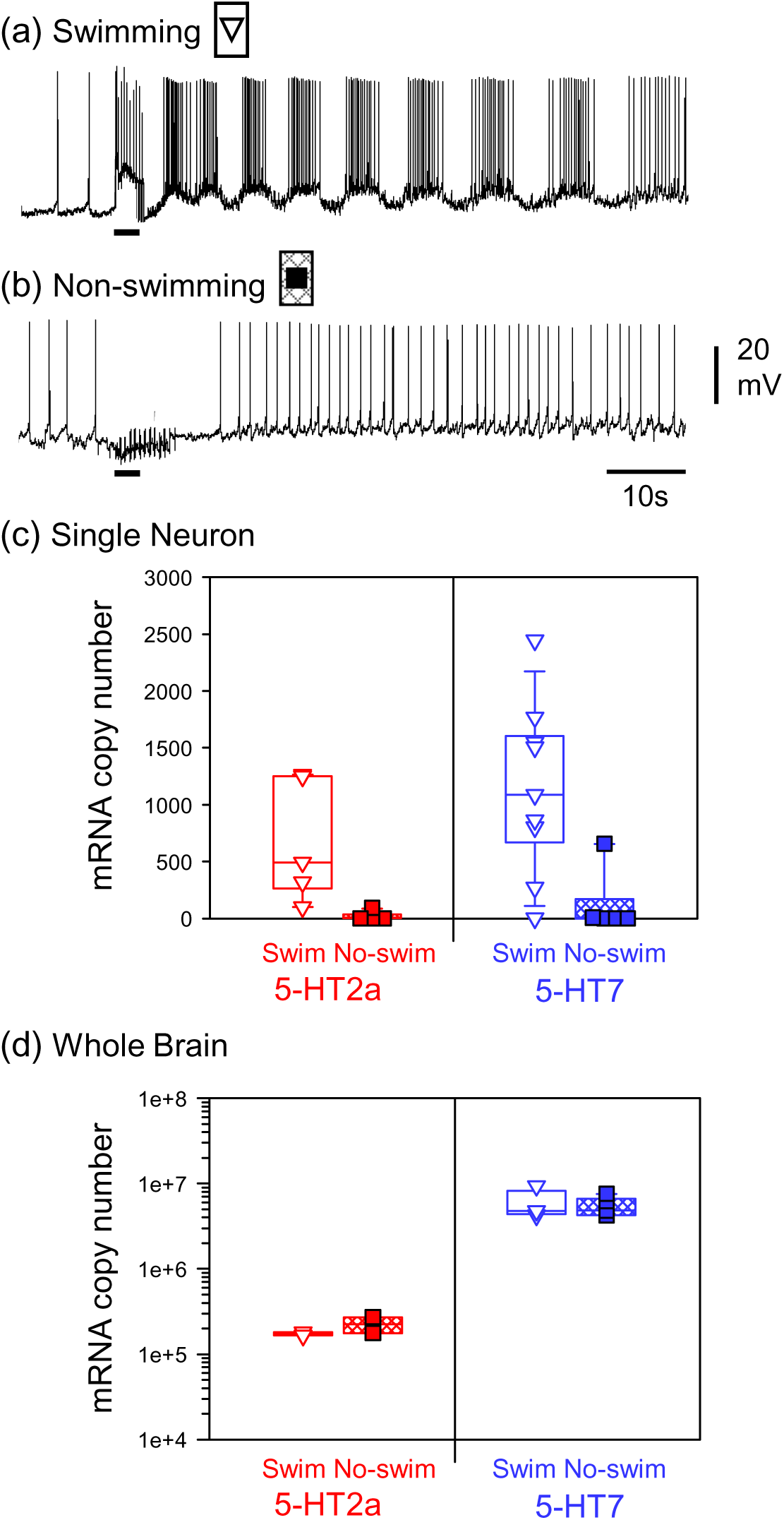
Variation in swimming and non-swimming *Pleurobranchaea*. Intracellular recordings of C2 neurons in response to body wall nerve stimulation (black bar) in swimming (a) and non-swimming (b) *Pleurobranchaea*. (c) Single neuron absolute qPCR measurements of mRNA copy number for 5-HT2a and 5-HT7 receptor genes in *Pleurobranchaea* C2 neurons from swimming (open triangles, white box) and non-swimming (filled squares, hatched box) preparations. Symbols represent results from individual C2 neurons. The boxes represent the range from 25% to 75% of sample expression levels. 95% confidence range is represented by error bars. (d) Whole brain absolute qPCR shows that there is no difference in expression of receptors between swimming and non-swimming preparations in total brain mRNA levels.

5-HT2a and 5-HT7 gene sequences were found in the transcriptome of C2s isolated from *Pleurobranchaea* that swam, but not in non-swimming specimens (Table 2). These results were validated by qPCR (Fig. 3C). Only one sample from the non-swimming *Pleurobranchaea* group showed low expression of 5-HT2a according to qPCR, calculated at 45 mRNA copies. There was a large degree of variability in the amount of 5-HT2a expressed in individual C2 samples from swimming individuals, from 100 to 1276 copies. All but one sample from a swimming preparation expressed 5-HT7 genes according to qPCR. Variability in qPCR-measured expression was highest for this gene, with a range of 0 to 2446. Only one of the seven C2 samples from non-swimming preparations exhibited 5-HT7 gene expression. This difference in expression was not seen in the total brain mRNA (Fig. 3D). Thus, *Pleurobranchaea* individuals that exhibited a swim motor pattern on the day of testing resembled *Tritonia* in the expression of 5-HT2a and 5-HT7 receptor genes in C2 neurons, whereas the C2 neurons of non-swimmers resembled *Hermissenda* in the absence of this expression.

**Table 2:** Comp IDs 5-HT receptors in *Pleurobranchaea* C2 transcriptome assemblies

| **Receptor** | **Swimming** | **Non-swimming** |
| --- | --- | --- |
| 5-HT1a | not found | not found |
| 5-HT1b | DN12981_c0_g1_i1 | not found |
| 5-HT2a | DN7433_c0_g2_i1 | not found |
| 5-HT2b | not found | not found |
| 5-HT4 | not found | not found |
| 5-HT6 | not found | not found |
| 5-HT7 | DN6728_c0_g1_i1 | not found |

We did not perform qPCR on the remaining receptor types, but relied on the transcriptomes. The swimming *Pleurobranchaea* C2 transcriptome contained sequences for 5-HT1b expression but none of the other receptors whereas the C2 transcriptome of non-swimmers did not contain any 5-HT receptor gene sequences (Table 2). This suggests that on the days that an individual *Pleurobranchaea* does not swim when tested, its C2 neuron did not express any 5-HT receptor genes, but it expressed 5-HT1b, 5-HT2a, and 5-HT7 receptor genes on days when it did swim.

### Expression of other G-protein coupled receptors did not correlate with swimming

We examined the single neuron transcriptomes to determine whether other GPCRs were differentially expressed in C2 of *Tritonia*, *Hermissenda*, and swimming and non-swimming *Pleurobranchaea* (Table 3). Molluscs express three identified dopamine receptor subtypes, known as D1, D2, and DInv [33]. The D1 dopamine receptor was found in the *Hermissenda* C2 but absent in the other samples, whereas the Dinv dopamine receptor was found only in the *Tritonia* C2 transcriptome and was missing in the others. In contrast, the D2 dopamine receptor was found in all four samples.

**Table 3:** Comp IDs of other GPCRs in C2 Transcriptome Assemblies.

|  | *Tritonia* | *Hermisenda* | *Pleur -Swim* | *Pleur – Non Swim* |
| --- | --- | --- | --- | --- |
| D1 Receptor | not found | DN56948_c0_g1_i1 | not found | not found |
| D2 Receptor | DN2095_c0_g1_i1 | DN66395_c0_g1_i5 | DN9070_c0_g1_i1 | DN2245_c0_g1_i1 |
| Dinv Receptor | DN36822_c0_g1_i1 | not found | not found | not found |
| Octopamine Receptor 1 | DN282_c0_g3_i1 | DN80161_c8_g1_i2 | not found | not found |
| Octopamine Receptor 2 | DN46207_c10_g1_i1 | DN61974_c0_g1_i1 | DN32740_c0_g1_i1 | not found |
| Adrenergic Receptor | DN28513_c0_g1_i1 | DN68537_c0_g2_i1 | not found | DN16230_c0_g1_i1 |
| mAChR | DN44275_c2_g1_i1 | DN81430_c5_g1_i1 | not found | not found |
| Histamine H1 Receptor | DN39405_c0_g1_i1 | DN81430_c5_g1_i1 | not found | not found |
| Neuropeptide Y receptor | DN46118_c9_g2_i3 | DN73680_c1_g3_i2 | not found | not found |
| Prolactin releasing receptor | DN45282_c6_g3_i1 | DN77362_c1_g2_i1 | not found | not found |
| Somatostatin Receptor | DN19522_c0_g1_i1 | DN82721_c1_g1_i4 | not found | not found |
| Orexin Receptor 1 | not found | DN76532_c0_g1_i1 | DN2724_c1_g1_i1 | DN2748_c0_g1_i1 |
| Orexin Receptor 2 | not found | DN45196_c1_g1_i1 | not found | not found |
| GPCR83 | not found | DN61803_c0_g1_i1 | not found | not found |
| GPCR161 | DN42626_c1_g1_i2 | not found | DN36378_c0_g1_i1 | not found |
| Cardioacceleratory Peptide Receptor | not found | DN79207_c0_g1_i1 | not found | not found |
| Neuromedin U Receptor | DN69884_c0_g1_i1 | DN73225_c0_g2_i1 | not found | not found |

Several other types of GPCR were found in both swimmers and non-swimmers. These GPCRs were also identified in the C2 transcriptomes from *Tritonia* and *Hermissenda*. (Table 3). However, the presence or absence of these other GPCRs did not correlate with swimming. A greater number of GPCR types were found in the two nudibranchs, compared with the *Pleurobranchaea* data sets. This is not due to an absence of these genes in the brain transcriptome, but an absence in the C2 transcriptome.

### Neuropeptide expression by C2 was consistent

C2 neurons are immunoreactive for SCP across nudipleura species [14]. The SCP precursor gene was found in C2 neuron transcriptomes from all three species in this study. Of note, there are two SCP precursor gene alternative splice isoforms present in molluscan brains, a short and long isoform [24]. In C2 neuron transcriptomes, only the short isoform was present (Supplemental Figure 1). Single-neuron qPCR confirmed that the SCP precursor gene is expressed in the isolated C2 neurons from each species, averaging 442 copies in *Tritonia*, 625 copies in *Hermissenda*, 989 copies in swimming *Pleurobranchaea*, and 197 copies in non-swimming *Pleurobranchaea*.

## Discussion

We found that differences in expression of particular 5-HT receptor genes in an identified neuron correlate with the presence of a type of swimming behavior. Using a combination of single-neuron transcriptome sequencing and absolute qPCR, gene expression for 5-HT receptors was measured in C2 neurons from three species of nudipleura sea slugs. The nudibranchs, *Tritonia* and *Hermissenda* differ in swimming behavior and in the expression of particular 5-HT receptor subtypes. *Pleurobranchaea* exhibits variations in swimming behavior and coordinate variations in the expression of 5-HT receptor genes; C2 neurons extracted from swimming individuals shared a profile of 5-HT receptor subtype expression with *Tritonia*, whereas non-swimming *Pleurobranchaea* individuals did not express any 5-HT receptor genes in this neuron. Overall, these results indicate a role for neuromodulatory gene expression in species-specific behaviors, as well as in individual variability in behavior.

### C2 synaptic modulation may be controlled via specific 5-HT receptors

In *Tritonia* C2 neurons, two subtypes were consistently measured across samples and showed expression above 200 copies per cell with single-neuron qPCR: 5-HT2a and 5-HT7. C2 neurons isolated from swimming *Pleurobranchaea* also expressed 5-HT2a and 5-HT7 subtypes. In contrast, C2 neurons from non-swimming *Pleurobranchaea* and *Hermissenda* did not express these two genes. Thus, the expression of 5-HT2a and 5-HT7 receptor genes correlated with the presence of the swimming behavior and the modulation of C2 synaptic strength by serotonin. In contrast, *Hermissenda* C2 neurons expressed 5-HT4 and 6, which were not found in *Tritonia* or *Pleurobranchaea* C2 neurons. The functions of these 5-HT receptors in *Hermissenda* C2 neurons are currently unknown.

Methysergide, a mammalian 5-HT1/2 family antagonist, blocks swimming in both *Tritonia* and *Pleurobranchaea* [18, 20]. Additionally, it reduces DSI-mediated modulation of C2 when bath applied to isolated *Tritonia* or *Pleurobranchaea* brain preparations [20, 22]. It is possible that methysergide acts on 5-HT2a receptors in *Tritonia* and *Pleurobranchaea* C2 neurons, consistent with its actions on 5-HT2 family receptors in mammals. However, methysergide binding selectivity may differ between receptor families in different animal phyla, as has been reported previously with respect to receptor pharmacology [34, 35].

### Other serotonin receptors may mediate additional responses

DSI stimulation and serotonin application cause three additional effects on C2 beside synaptic enhancement: a decrease in spike frequency adaptation that is associated with a decrease in C2 spike after-hyperpolarization, a fast depolarizing potential, and a slow depolarization [22, 23]. The slow depolarization is blocked by methysergide, suggesting that it may also be mediated by 5-HT2a receptors. The fast potential is blocked by the antagonist gramine, which does not block the synaptic enhancement or the slow potential, suggesting that it is mediated by a different 5-HT receptor. The “fast” potential is still fairly slow by synaptic standards with a decay time constant of 0.28 seconds, compared to a decay time constant of 4.4 seconds for the slow potential, indicating that both could be mediated by a GPCR.

C2 reliably expressed 5-HT1b receptor genes in both *Tritonia* and *Hermissenda,* and this sequence was present in the transcriptome of swimming *Pleurobranchaea*. However, this gene was expressed at fewer than 200 copies per cell and thus was not as prevalent as the other 5-HT receptor genes. Because it was expressed in both *Tritonia* and *Hermissenda*, the 5-HT1b receptor may mediate an action of serotonin on C2 other than enhancement of synaptic release or synaptic potentials, none of which were seen in *Hermissenda*. The effect of serotonin on spike frequency adaptation was not tested in *Hermissenda* or *Pleurobranchaea*, leaving open the possibility that it is mediated by 5-HT1b receptors.

The expression of multiple neuromodulatory receptors is a feature that has been observed in several single-neuron studies. In the anaspid mollusc *Aplysia californica*, pleural ganglion sensory neurons express at least three 5-HT receptor subtypes, identified with single cell PCR [33]. In crustaceans, individual neurons respond to a variety of neuromodulatory substances [36, 37].

### Other genes did not show a correlation with behavior

Several GPCRs were identified in C2 neuronal transcriptomes from each species, indicating that C2 may have several neuromodulatory inputs. Furthermore, many these receptors exhibited differed in their expression across species, but did not correlate with the swimming behavior. In the case of *Pleurobranchaea*, it is worth noting that the sample size for the transcriptome sequencing was lower than that of the other two species (see methods). Therefore, it is possible that *Pleurobranchaea* C2 neurons express more GPCRs than those captured by sequencing in this study. There may be other, as yet unidentified, species-specific behaviors mediated by C2 neurons.

Although the expression profile of 5-HT receptors in C2 was species-specific and variable across individuals with different behaviors, the SCP precursor gene was expressed in all C2 neurons, regardless of species and behavior. SCP immunoreactivity was previously identified as a defining characteristic of C2 neurons and an indication of their homology across species [14]. The presence of SCP mRNA in C2 neurons from each species confirmed the identity of the cell, and showed that there are gene markers found across homologous neurons, regardless of species. However, the lower expression of SCP in the C2 neurons of non-swimming *Pleurobranchaea* suggests either an overall reduction in gene expression or the presence of a gene expression network which co-regulates the expression of SCP and the 5HT 2a/7 receptors.

### Parallel evolution of neuromodulation

Most nudipleura species do not exhibit DV swimming. Furthermore, based on their phylogenetic tree, it appears that *Tritonia* and *Pleurobranchaea* do not share a most recent common ancestor that exhibited this behavior and thus likely evolved DV swimming independently [38]. Both species also gained serotonergic modulation of C2 neurons independently [20]. The expression of orthologous 5-HT2a and 5-HT7 receptor genes in C2 neurons from swimming *Pleurobranchaea* and *Tritonia* suggests that the similar neuromodulatory responses in these two species are a result of parallel evolution, where homologous genes came to be used independently for the same function [39]. It was previously proposed that serotonergic modulation transformed C2 and other neurons from a latent circuit into the functional swim motor pattern CPG circuit observed in *Tritonia* and *Pleurobranchaea* [20]. In fact, it may be that the presence of 5-HT2a and/or 5-HT7 is the “switch” in this evolutionary transformation.

Although the mechanism by which the 5-HT receptor subtypes affect behaviors involving C2 is currently unknown, the correlation between their expression and swimming behaviors shown here presents a tantalizing possibility that manipulating 5-HT2a or 5-HT7 expression in non-swimmer C2 neurons might cause DV swimming to occur, while knocking down their expression in DV-swimmers may reduce or eliminate the swim motor pattern.

This type of experiment was performed in prairie and montane voles. The neuromodulatory vasopressin receptor V1a is expressed by neurons in the ventral pallidum of monogamous prairie voles, but not in that region in the closely related, non-monogamous montane voles [40]. Virally-induced V1a receptor expression in the ventral pallidum of the the non-monogamous montane voles causes them to exhibit behavior characteristic of monogamy [41]. Thus, the presence or absence of the V1a receptor is the underlying mechanism that facilitates the behavioral switch.

### C2 showed within-species variability in amount of 5-HT receptor mRNA expressed

There was variability in the level of expression of some receptor subtypes within each species as measured by single-neuron qPCR. 5-HT7 was expressed in one of six *Hermissenda* C2 samples and in one non-swimming *Pleurobranchaea* sample. 5-HT1a was expressed in one *Tritonia* C2 neuron sample, but it was below the level of qPCR detection in the other samples tested for that gene. Similarly, 5-HT2b was expressed in only a few of the *Tritonia* and *Hermissenda* C2 neurons tested. In the remaining qPCR trials, there was variability in the amounts of each subtype expressed between samples from within a species.

This variability could be due to natural fluctuations in the amount of mRNA for a given gene, which occurs randomly in many cell types [42, 43]. In all eukaryotic cells, mRNA molecules are transcribed in the nucleus, translated to proteins by ribosomes, and then degraded as part of normal cell cycling. It is therefore to be expected that mRNA amounts would wax and wane over time, causing variability in qPCR measurements. It is possible that 5-HT receptor mRNA fluctuates randomly in this way.

In other systems, 5-HT receptor mRNA expression variability correlates with specific behaviors. In the neural circuit controlling song production in birds, microRNAs alter seasonal expression of 5-HT and other receptors [44]. Estrogen produced during reproductive phases causes a reduction in 5-HT receptor mRNA in mussel gonads [45].

In non-human primates and rats, 5-HT receptors and other 5-HT-related genes vary with stress in several brain regions [46–49]. 5-HT receptor mRNA was found to vary with circadian rhythm and hibernation periods in rodents [50, 51]. The fluctuation of 5-HT receptor mRNA in C2 neurons may have as yet undiscovered functional consequences. It is also worth noting that the animals used in this study were wild-caught adults. They may have exhibited varying mRNA expression because of their individual experiences in the wild before ending up in the laboratory.

### Non-swimming Pleurobranchaea did not express any 5-HT receptors in C2

Swimming varies daily in *Pleurobranchaea*, as measured using both *in vivo* behavioral assays and *in vitro* fictive swimming and this variability correlates with serotonergic enhancement of C2 synaptic strength [20]. Here, we found that for the most part, C2 in non-swimming *Pleurobranchaea* did not express any 5-HT receptor genes. Although the presence of mRNA in a cell does not necessarily indicate that the corresponding protein is present [52], the correlation of 5-HT receptor mRNA expression and swim motor pattern bursts by C2 in *Pleurobranchaea* points to a genetic mechanism that may at least partially facilitate the mechanism by which C2 and other neural components become functional as the swim motor pattern circuit. *Pleurobranchaea* C2 samples exhibited the most variability by qPCR for 5-HT2a, 5-HT7, and SCP. Perhaps there is a species-specific aspect of mRNA cycling or co-expression that causes this increased variability. If this were the case, then over time there could be periods where few 5-HT receptor proteins were expressed at C2 synapses in *Pleurobranchaea*, which could cause temporary loss of DV swimming and serotonergic modulation.

---

**Supplemental Table 1:** *Pleurobranchaea* whole-brain transcriptome metrics.

| ***Pleurobranchaea***<br>**Whole Brain Transcriptome** |  |
| --- | --- |
| **Raw Illumina data** |  |
| No. of reads (2x100) | 106,288,872 |
| % Phred Score >30 | 91.3% |
| **Assembled Components** |  |
| Trinity transcripts | 217,916 |
| Trinity genes | 147,024 |
| %GC | 40.4% |
| N<sub>50</sub> | 2,102 |
| Median length | 360 |
| Average length | 902 |
| Total bases | 196,485,559 |
| Longest contig | 5,008 |

**Supplemental Table 2:**
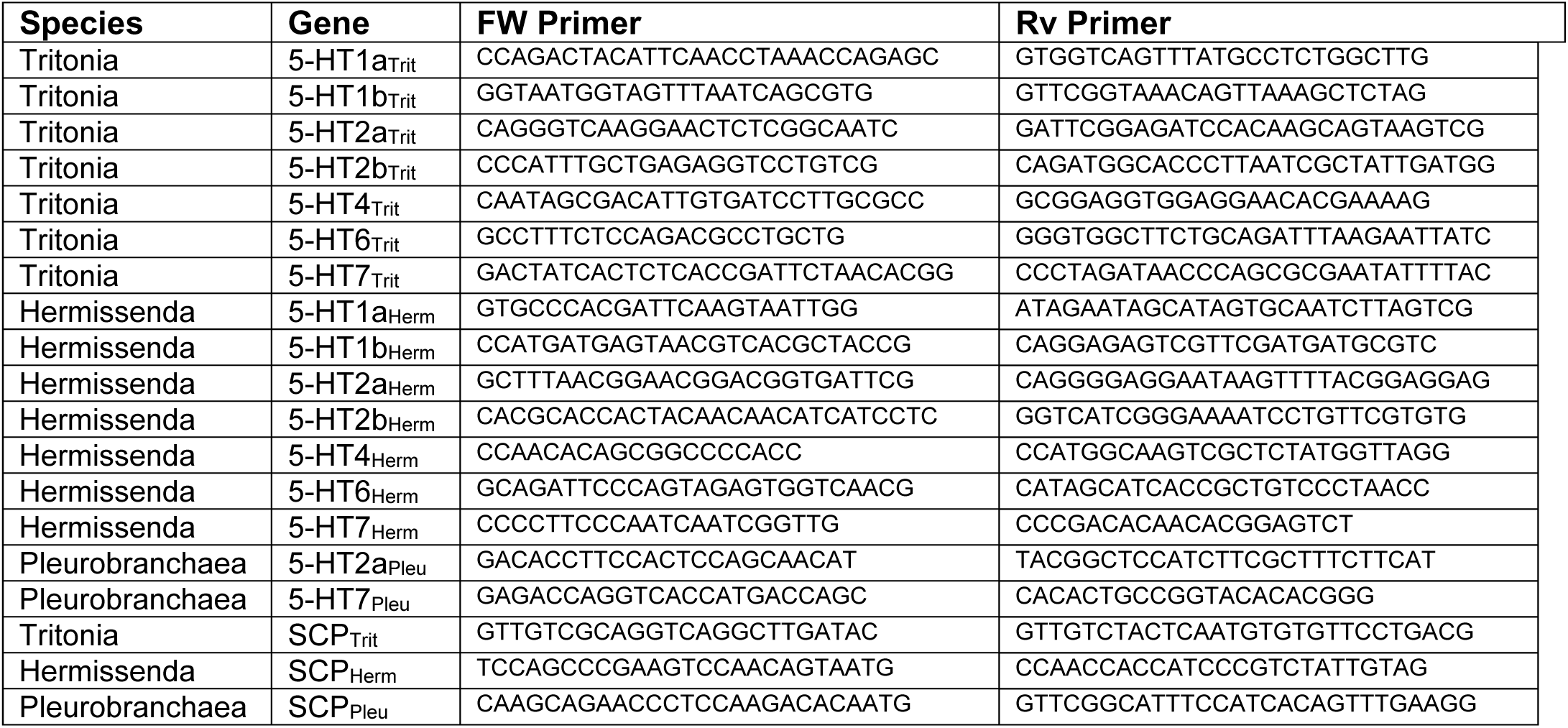
Primers used for initial amplification and plasmid cloning.

**Supplemental Table 3:**
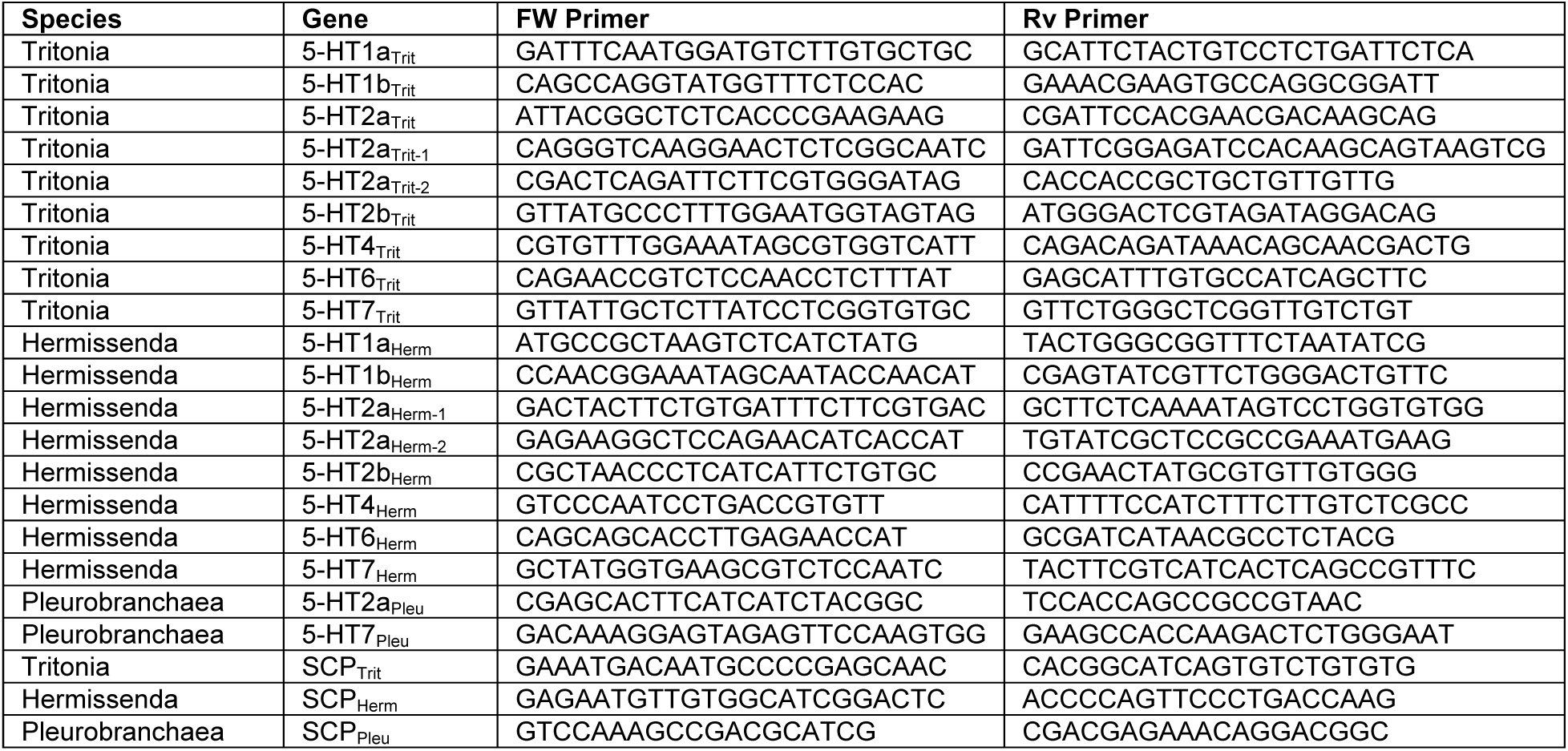
Primers used for absolute qPCR.

**Supplemental Table 4:** *Pleurobranchaea* whole brain transcriptome component identification numbers.

| Gene | Comp ID |
| --- | --- |
| 5-HT1a<sub>Pleu</sub> | comp77412_c0_seq1 |
| 5-HT1b<sub>Pleu</sub> | comp71667_c0_seq4 |
| 5-HT2a<sub>Pleu</sub> | comp68325_c1_seq1 and comp70123_c2_seq3 |
| 5-HT2b<sub>Pleu</sub> | comp53876_c0_seq1 |
| 5-HT4<sub>Pleu</sub> | comp159323_c0_seq1 and comp48526_c1_seq1 |
| 5-HT6<sub>Pleu</sub> | comp68387_c1_seq1 |
| 5-HT7<sub>Pleu</sub> | comp73047_c11_seq1 |

**Supplemental Figure 1:**
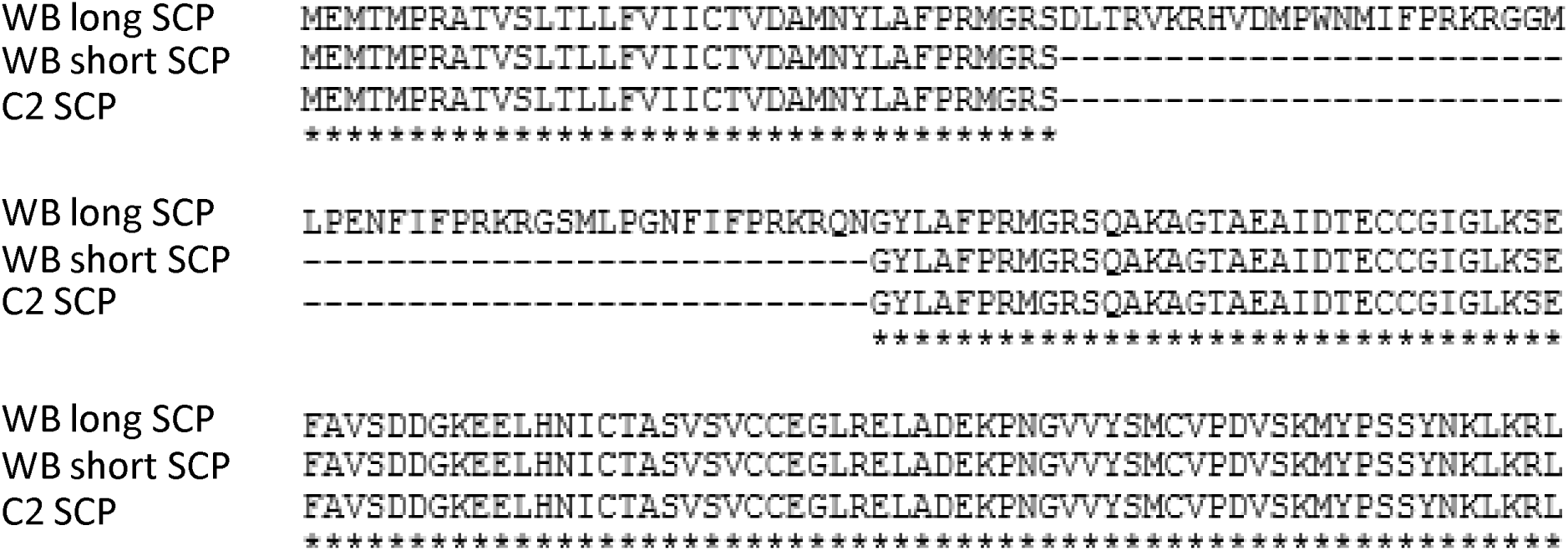
SCP alignment from whole brain and C2 transcriptomes. *Tritonia* whole brain (WB) short and long SCP sequences were aligned with the corresponding C2 neuron transcriptome sequence.

